# Age and genetic background determine hybrid male sterility in house mice

**DOI:** 10.1101/382382

**Authors:** Samuel J. Widmayer, David L. Aylor

## Abstract

Hybrid male sterility (HMS) is a unique type of reproductive isolation commonly observed between house mouse *(Mus musculus)* subspecies in the wild and in laboratory crosses. We identified hybrids that display three distinct trajectories of fertility despite having identical genotypes at the major HMS gene *Prdm9* and the X Chromosome. In each case, we crossed female PWK/PhJ mice representative of the *M.m.musculus* subspecies to males from classical inbred strains representative of *M.m.domesticus:* 129S1/SvImJ, A/J, C57BL/6J, and DBA/2J. PWK129S1 males are always sterile, while PWKDBA2 males escape HMS. In addition, we observe age-dependent sterility in PWKB6 and PWKAJ males. These males are fertile between 15 and 35 weeks with moderate penetrance. These results point to multiple segregating HMS modifier alleles, some of which have an age-dependent mode of action. Age-dependent mechanisms could have broad implications for the maintenance of reproductive barriers in nature.

**Author Summary:** Two subspecies of house mice show partial reproductive barriers in nature, and may be in the process of speciation. We used mice derived from each subspecies to replicate hybrid male sterility (HMS) in laboratory mice. Two major genetic factors are well established as playing a role in mouse HMS, but the number of additional factors and their mechanisms are unknown. We characterized reproductive trait variation in a set of hybrid male mice that were specifically designed to eliminate the effects of known genetic factors. We discovered that age played an important role in fertility of some hybrids. These hybrid males showed a delayed onset of fertility, then became fertile for only a few weeks. Across all hybrids males in our study, we observed three distinct trajectories of fertility: complete fertility, complete sterility, and age-dependent fertility. These results point to two or more critical HMS variants with large enough effects to completely restore fertility. This study advances our understanding of the genetic architecture and biological mechanisms of reproductive isolation in mice.

## Introduction

Hybrid male sterility (HMS) is a special type of reproductive isolation wherein crosses between genetically distinct groups produce viable, yet sterile male offspring. The Dobzhansky-Muller model of reproductive isolation [1–3] proposes an evolutionary genetic mechanism for the development of reproductive incompatibilities. With enough restriction to gene flow, diverging populations accumulate and fix new mutations. While these unique alleles are neutral within each population, these alleles act deleteriously in hybrids through epistatic interactions that cause HMS.

House mice *(Mus musculus)* are a powerful system for studying HMS. House mice have a cosmopolitan distribution and exist in three genetically distinct subspecies: *M. m. musculus, domesticus,* and *castaneus* [4–6]. These subspecies began diverging approximately 500 thousand years ago [7], yet gene flow between the subspecies is still substantial. A narrow hybrid zone exists between the *musculus* and *domesticus* subspecies, which have natural distributions across eastern and western Europe, respectively. These two subspecies are in the earliest stages of reproductive isolation. Allele frequencies exhibit sharp clines across the hybrid zone [8–10], and reduced fertility is common in mice with relatively high degrees of subspecies admixture [11]. These findings provide strong support for the reduction of gene flow between subspecies due to partial reproductive isolation in wild mice.

Studies of natural mouse populations have revealed few candidate HMS loci, and others have made progress by crossing inbred mouse strains representative of the major mouse subspecies [12]. The majority of classical inbred mouse strains are are genetic mosaics of the subspecies but the vast majority of the genome consists of *domesticus* ancestry [13]. Hybrid sterility is generally asymmetric in crosses between primarily *domesticus-* and musculus-derived inbred strains, affecting only hybrid males derived from *musculus* dams and *domesticus* sires [14–17]. Only one gene and one QTL have been definitively linked to the development of HMS. *Prdm9* is a histone methyltransferase that in necessary for the formation of the synaptonemal complex [15–20]. *Hstx2* [21–23] is a QTL on Chr *X^Msc^* that is necessary for HMS in all reported *musculus* x *domesticus* hybrids. Specific genotypes at these loci lead to asynapsis during pachytenesis [15, 16, 18] and to impaired meiotic sex chromosome inactivation (MSCI) [17, 21, 24–26] and subsequently to HMS.

Allelic variation at *Prdm9* has been shown to play a key role in most studies of mouse HMS[15]. In fertile mice, *Prdm9* binds DNA and demarcates recombination hotspots by directing double-stranded break (DSB) initiation sites immediately preceding the synapsis of homologous chromosomes [27, 28] during spermatogenesis. The *Prdm9^Dom2^* allele exhibits DNA-binding motif variation relative to *Prdm9^Msc^* or *Prdm9^Dom3^,* and the protein isoforms exhibit allele-specific binding genome-wide [29–32]. Aberrant DNA-binding in *Prdm9* heterozygotes results in asynapsis [15, 16, 19, 27, 29, 33]. Fertility in this system can be rescued by decreasing the degree of asymmetric Prdm9-binding through transgenic rescue [16], replacing the *Dom2* allele with *Dom3* or humanized *Prdm9* alleles) [16, 18, 19], or by artificially creating symmetric homologs [34]. One model of HMS in mice posits that the exact locations of the asymmetric hotspots matter less than their abundance or density. In this model, *Prdm9* is the only essential HMS gene. Diffuse genetic background effects are not dependent on specific HMS alleles, since almost any stretch of asymmetric double-strand break repair could be the cause of asynapsis. The key prediction of this model is that the proportion of *domesticus* ancestry should predict the degree of aberrant *Prdm9* binding and subsequently the degree of asynapsis.

Although *Prdm9* and *Hstx2* are important drivers of HMS, they are not sufficient to do so on all genetic backgrounds. Wild mice captured from the hybrid zone are frequently fertile, indicating that specific genetic architectures rescue fertility [11, 35]. Moreover, male progeny of C57BL/6J-Chr 17^PWD^ consomic sire and C57BL/6J-Chr X^PWD^ consomic dam carry the known HMS genotypes and are fertile, indicating the existence of additional alleles that can rescue fertility in the B6 background [22]. *Prdm9* has at least two segregating alleles within the *domesticus* subspecies. Intersubspecific hybrid male mice that carry *Prdm9^Dom2^* (e.g. C57BL/6J) are generally sterile, while hybrid male mice that carry *Prdm9^Dom3^* (e.g, WSb/EiJ) are typically fertile. However, reproductive phenotypes segregated in F2 intercrosses derived from WSB, and several QTL have been associated independent of *Prdm9* or Chr X [35–37]. Other forward genetics approaches have identified large-effect QTL in the wild [35] and between musculus-derived inbred mouse strains [14, 38]. Thus, while *Prdm9* and *Hstx2* explain a large proportion of variation in HMS phenotypes, the complete genetic architecture of HMS is unknown.

We found that PWK-derived F1 hybrids displayed three distinct trajectories of HMS that were dependent on age and genetic background: complete sterility throughout life, complete fertility throughout life and age-dependent HMS. This finding is significant because these hybrids all harbored identical genotypes at the two major HMS loci, *Prdm9* and Chr X. Therefore, HMS is necessarily linked to undiscovered alleles segregating between these strains. We measured fertility profiles and the cellular phenotypes that were linked to HMS at three ages for each hybrid. Lastly, we identified regions of subspecific ancestry that are candidates to harbor HMS alleles. Together, these results provide the first evidence for age-dependent HMS in the mouse and substantially advance our understanding of genetic reproductive isolation in mice.

## Results

### Genetic background controls HMS phenotypic variation

We crossed PWK females to males of four different inbred mouse strains: 129S1, A/J, B6, and DBA2 (**Figure 1A**) and measured reproductive phenotypes in the resulting focal hybrid males at 8 weeks of age. Males across this panel had invariant *Prdm9* and Chr X genotypes, which allowed us to directly test the effects of background genetic variation on HMS traits. We also generated reciprocal hybrids by crossing 129S1, A/J, B6, and DBA2 females to PWK males (**Figure 1B**). These mice were similarly invariant at *Prdm9* but lacked the *musculus* Chr X that previously has been linked to HMS.

**Figure 1:**
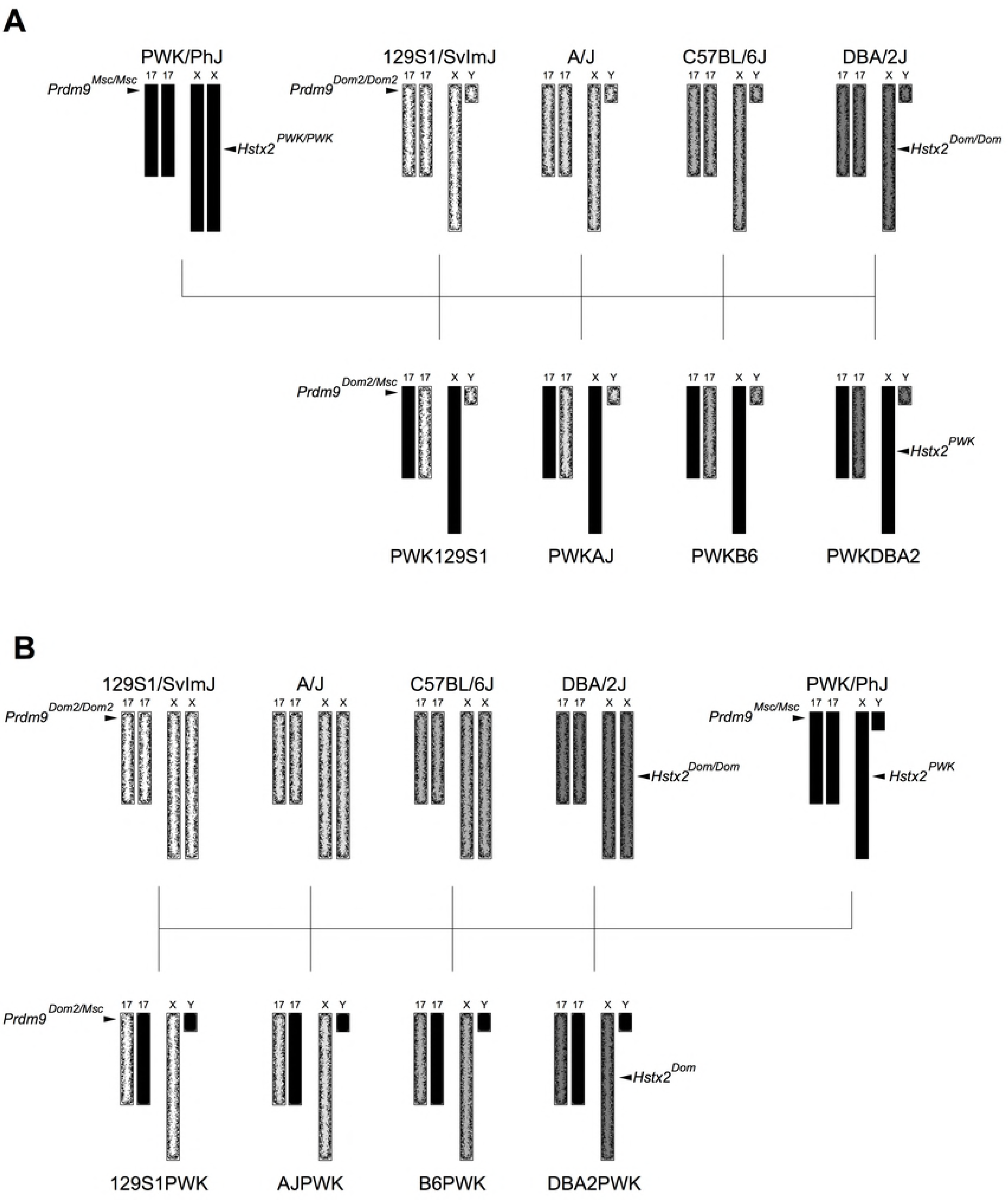
Crossing scheme to generate genetically diverse hybrid male mice. A) We generated focal hybrid males by crossing PWK (black) females to males of four classical inbred mouse strains: 129S1, A/J, B6, and DBA2 (stippled). These four strains all carry the *Prdm9^Dom2^* allele that has been previously linked to HMS in other crosses. Focal mice are fixed for the *Prdm9^Dom2/Msc^* genotype and PWK Chr X, eliminating the effects of those major HMS loci. B) Reciprocal hybrids were generated by reversing the cross direction.

PWK129S1, PWKAJ, and PWKB6 mice displayed reproductive phenotypes consistent with sterility (**Figure 2, Table S1**). All three of these hybrid males had reduced combined testes weights (**Figure 2A**) and total sperm counts (**Figure 2B**) relative to their reciprocal hybrids (p ≤ 0.001). These results are important because both PWKAJ and PWKB6 have previously been reported to be fertile hybrid mice. In fact, fertile PWKAJ and fertile PWKB6 mice were necessary contributors to the two large mouse population-based resources, the Collaborative Cross (CC) and the Diversity Outbred (DO) population [39, 40]. In contrast, PWK129S1 males were known to be sterile [41]. In addition, PWKDBA2 males displayed substantially increased reproductive phenotypes relative to the other focal hybrids *(p ≤* 0.001). PWKDBA2 testes weights were indistinguishable from reciprocal DBA2PWK males (p = 0. 7813) yet had reduced sperm counts (p ≤ 0.0002) that were intermediate between reciprocals and the other focal hybrids.

**Figure 2:**
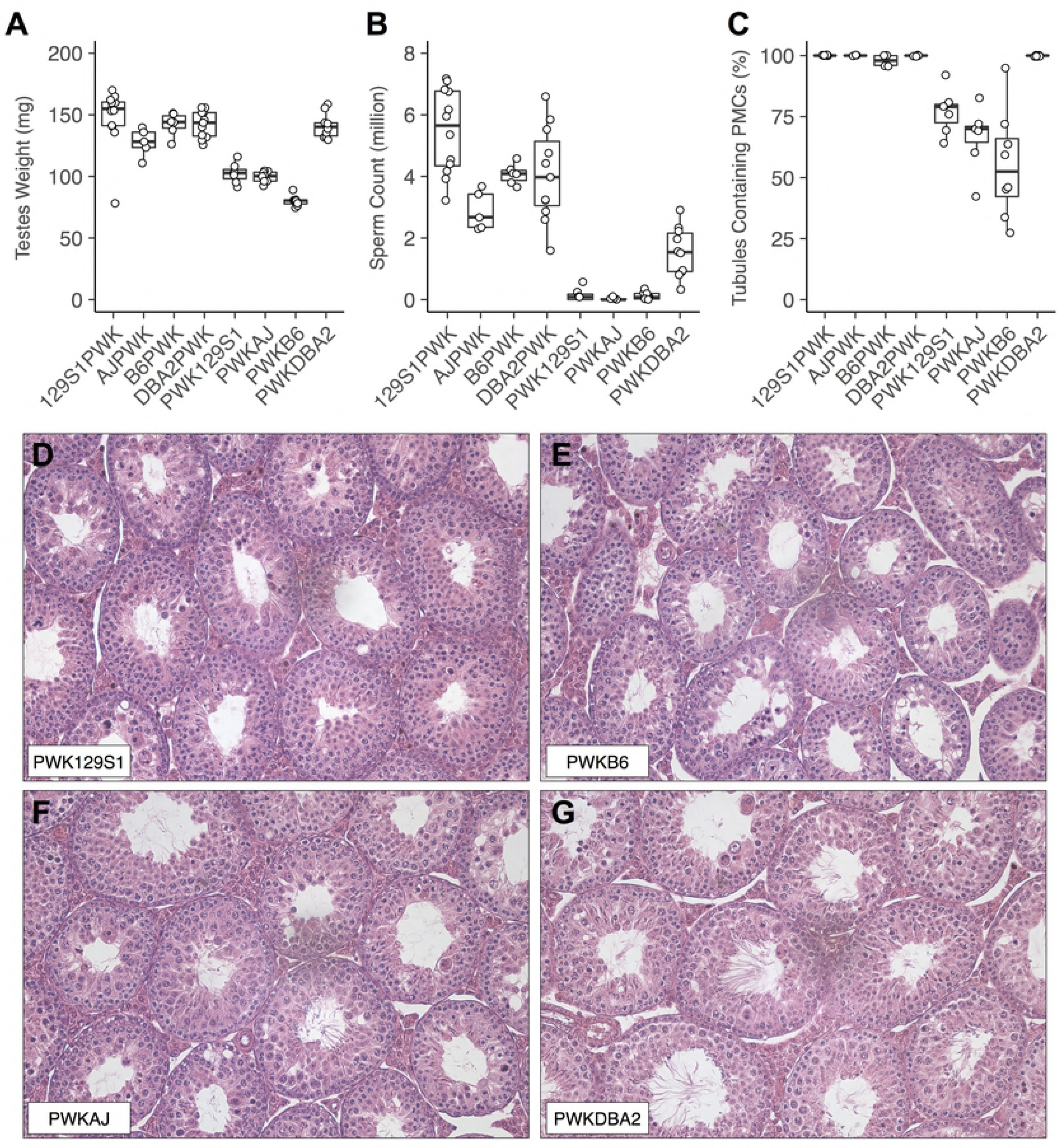
Reproductive phenotypes of 8-week old hybrid male mice vary across genetic backgrounds. A) Combined testes weight, B) sperm count, and C) percent of seminiferous tubules containing post-meiotic germ cells (PMCs) show significant variation between genetically distinct hybrids. Only PWKDBA2 hybrid males displayed phenotypes consistent with fertility at 8 weeks of age. D) Histological cross section of a representative PWK129S1, E) PWKB6, F) PWKAJ, and G) PWKDBA2 testis. PWKDBA2 seminiferous tubules contain abundant spermatids and spermatozoa, while many PWK129S1, PWKAJ, and PWKB6 tubules contain few to no spermatids, indicative of partial meiotic arrest.

These phenotypes suggested meiotic failure consistent with Prdm9-driven HMS. To test this, we analyzed histological cross-sections from the left testis of hybrid male mice, and estimated the percentage of seminiferous tubules within each cross-section that contained post-meiotic germ cells (PMCs) as a proxy for successful spermatogenesis (**Figure 2C**). We found that PWK129S1 (77%), PWKB6 (55%), and PWKAJ (67%) males each had a smaller fraction PMCs compared to PWKDBA2 males (**Figure 2D–2F**) (*p_adj_* ≤ 0.016) or their reciprocal hybrids (*p* ≤ 0.001). The percent of seminiferous tubules with PMCs for PWKDBA2 and all reciprocal hybrids was at least 96% (**Figure 2G**) with a median of 100%. These results showed that sterility in PWK129S1, PWKB6, and PWKAJ males was meiotic or pre-meiotic in mechanism. However, these PWK-derived hybrids did not display a complete meiotic block as has been shown in other intersubspecific hybrids [15, 16]. These findings also confirmed that spermatogenesis in PWKDBA2 hybrids is virtually unaffected.

These results clearly showed that one or more genetic variants in the DBA2 strain rescued fertility even in the presence of *Prdm9^Dom2^* and a PWK Chr X. In addition, these results introduced a conundrum. In contrast to our expectations, PWKAJ and PWKB6 males were sterile at 8 weeks of age and mirrored the expected sterility of PWK129S1 hybrids in both phenotype and cell composition.

### Fertility is age-dependent in PWKB6 and PWKAJ hybrids

Two separate observations suggested that age might help reconcile our initial results in PWKB6 hybrid males with their expected fertility. First, a colleague reported anecdotally that PWKWSB hybrids became prematurely sterile during an independent experiment investigating paternal age effects, often after successfully siring several litters (James Crowley, personal communication circa June 2011). We designed a breeding experiment to investigate this observation and to see if a similar phenomenon applied to PWKB6 males. Second, Forejt and colleagues reported that PWKB6 hybrid males exhibit delayed onset of fertility [18] during the course of our initial experiments. Therefore, we subsequently ran a second breeding experiment to characterize the onset of fertility. We crossed adult PWKWSB males (*n*=55) and PWKB6 males (*n*=63) that were at least 15 weeks of age to FVB females and continuously mated them until they ceased producing offspring. WSB harbors a *Prdm9^Dom3^* allele and PWKWSB hybrids are generally thought to be fertile. Given the importance of the *Prdm9^Dom2^* allele in PWKB6 HMS, we had no specific expectation of an age effect.

Only 40% of PWKB6 males sired offspring and the number of litters decreased with age. Furthermore, no PWKB6 males sired offspring after 35 weeks of age (**Figure 3**). PWKWSB males also showed a premature sterility, but at a much more advanced age. Nearly all PWKWSB males were fertile at 20 weeks of age (94%), but none were fertile after 58 weeks of age, with fewer than half of males siring litters. We concluded that HMS in PWKB6 hybrid male mice had a high but incomplete penetrance. In addition, we concluded that PWK-derived interspecific hybrids that are fertile during early life exhibited sterility past specific age points.

**Figure 3:**
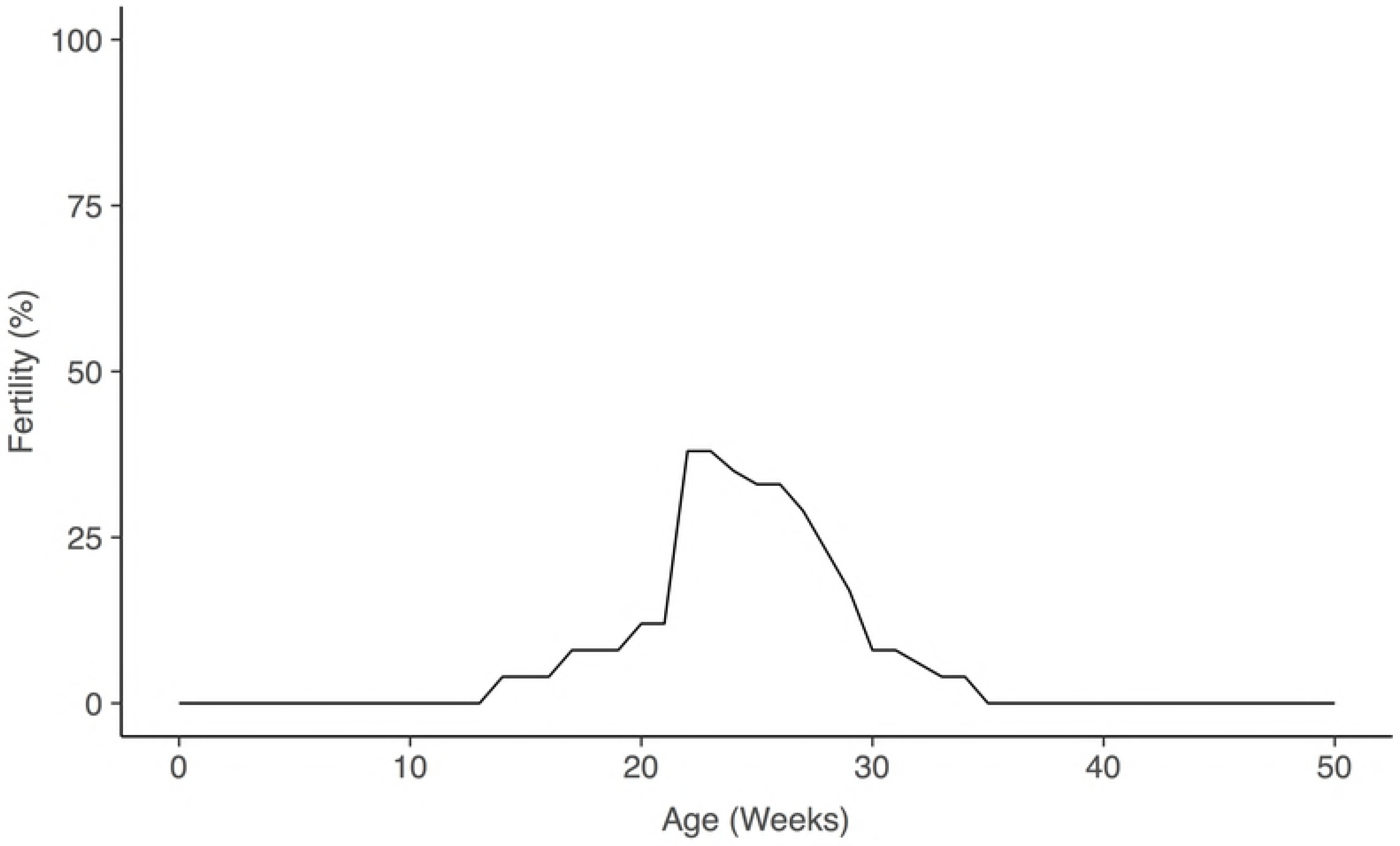
Inter-individual variation and age-dependence of fertility among PWKB6 hybrids. Fertility curve constructed from combining two breeding experiments. HMS had a ?60% penetrance, but some mice were fertile at ages 15-35 weeks. One experiment was designed to determine the age of fertility onset. The other found the cessation of fertility by age 35 weeks.

Moreover, this age-dependent sterility occurred in both *Prdm9^Dom2^* and *Prdm9^Dom3^* genetic backgrounds. These results established age as a critical factor in our system, but they did not explain why PWKB6 mice were sterile at age 8 weeks. If we had expected 60% of PWKB6 males to be sterile, our observation of 100% sterility was unlikely (p = 0.0168 in a *Binomial(8,0.6)* distribution).

To determine the age of fertility onset, we crossed young mice (5-8 weeks) to fertile females and measured latency until the first successful mating. PWKB6 males (*n*=17) bred continuously until age 20 weeks, and additional mice (*n*=9) were bred continuously until age 15 weeks. We also included PWKAJ males (*n*=6), reciprocal B6PWK males (*n*=6), and PWKDBA2 males (*n*=3). All PWKDBA2 and four of six B6PWK males sired offspring by 8 weeks of age, consistent with our initial screen. In contrast, the majority of PWKB6 and PWKAJ hybrid males were sterile. However, three of 26 PWKB6 males and two of six PWKAJ males sired litters at ages ranging from 12 to 20 weeks. All litters sired by PWKB6 consisted of only one pup, and the litters sired by PWKAJ males were of two and three pups. These litter sizes were substantially reduced compared to those sired by reciprocal B6PWK males (5-11 pups). These results supported our conclusion that sterility in PWKB6 male mice had a high but incomplete penetrance, and extended this result to PWKAJ. In addition, those PWKB6 and PWKAJ males that were fertile experienced a delay in the onset of fertility with respect to reciprocal hybrids, and showed reduced fertility based on litter sizes.

Having established the importance of age, we collected reproductive phenotypes of the focal hybrid males and their reciprocals at ages 20 weeks and 35 weeks (**Figure 4**). PWK129S1 males exhibited little change in testes weight or sperm counts, which remained under one million (p ≤ 0.99). In addition, PWK129S1 males displayed a reduction in the percentage of seminiferous tubules with PMCs between 8 and 20 weeks, from a mean of 77% at age 8 weeks to 52% at 20 weeks (p ≤ 0.0158). These results were consistent with complete sterility in PWK129S1 males.

**Figure 4:**
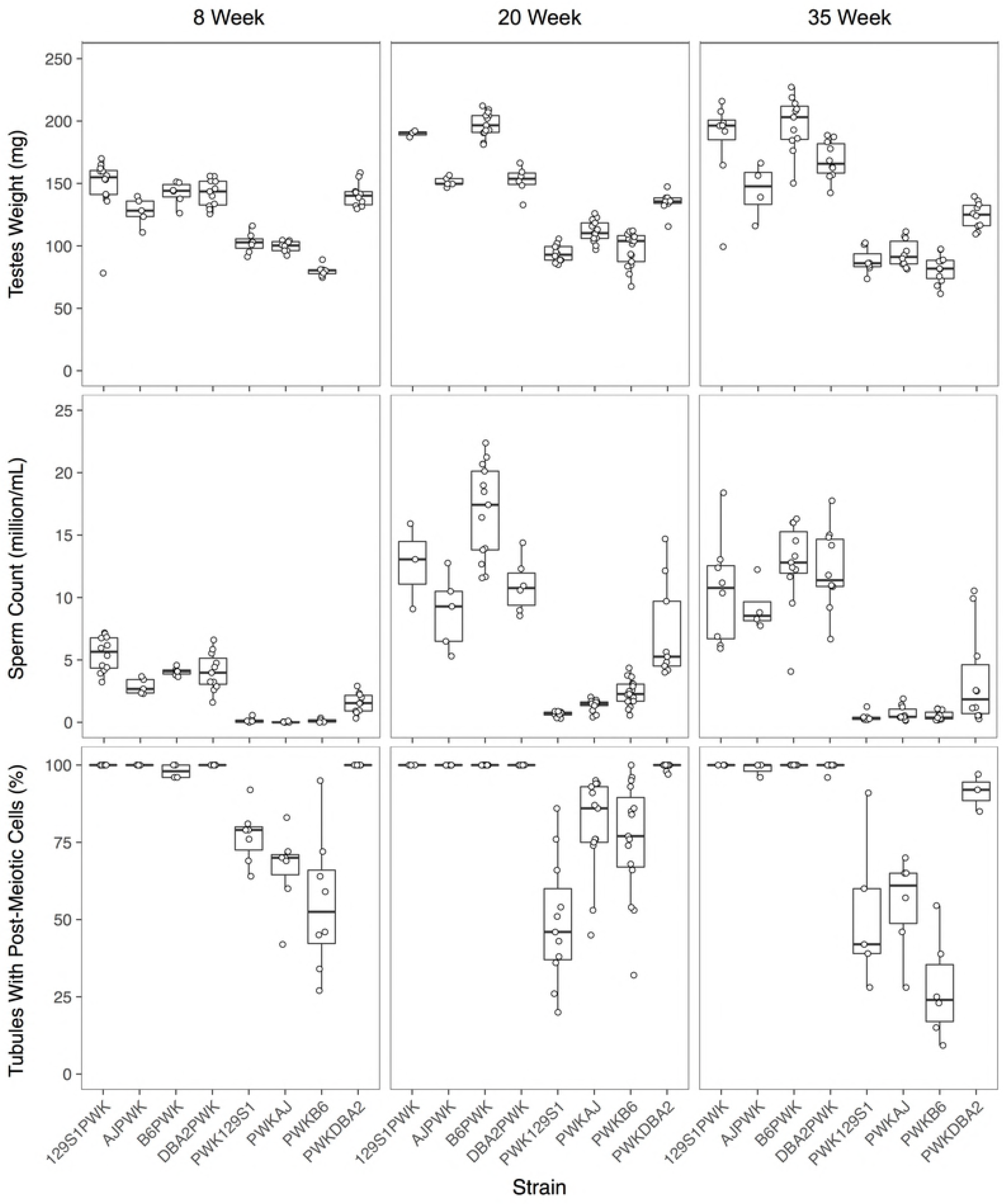
Genetic background dependent changes to HMS phenotypes over time. Testes weight, sperm count, and the percentage of seminiferous tubules containing post-meiotic germ cells (PMCs) across three ages: 8 weeks, 20 weeks, and 35 weeks of age.

In contrast, PWKDBA2 mice exhibited phenotypes consistent with fertility throughout life. Testes weight showed no change (p ≤ 0.3533), but sperm count increased by 20 weeks (p ≤ 0.0019) and then declined somewhat by 35 weeks (p ≤ 0.0423). Similar patterns were seen in the reciprocal hybrid males, though PWKDBA2 males never approached the high sperm counts of their reciprocals (p ≤ 0.001). The proportion of seminiferous tubules with PMCs exhibited no significant changes with age and PWKDBA2 males looked similar to reciprocals, except for a decline to 91% at 35 weeks (p ≤ 0.139). We concluded that age does not substantially impact the fertility of PWKDBA2 mice.

PWKB6 and PWKAJ mice exhibited marked increases in testes weight, sperm count, and the proportion of seminiferous tubules containing PMCs at age 20 weeks in comparison to 8 weeks (2-way ANOVA *p ≤* 0.001). The average PWKB6 combined testes weight increased by 22%, and the average sperm count displayed a 20-fold increase. The average PWKAJ combined testes weight increased by 12%, and sperm count increased over 60-fold. There was substantial variation in sperm count relative to 8 weeks in both PWKB6 (561,000 to 4.4 million) and PWKAJ (418,000 to 2 million). The proportion of seminiferous tubules containing PMCs ranged from 32% to 100%. These high variances were consistent with incomplete penetrance of HMS in PWKB6 and PWKAJ hybrids. All three fertility traits declined between 20 and 35 weeks of age in both PWKB6 and PWKAJ and were consistent with sterility. 94% of PWKWSB males were fertile throughout the 20-35 weeks window, indicating no effects until much later in life.

These complementary experiments gave us the first clear picture of a complex fertility curve in PWKB6 and PWKAJ male mice. Most of these hybrid males were sterile. For others, fertility was transient. The onset of fertility was delayed until age 12 weeks or later. Even after siring one or more litters, these hybrid males were all sterile by age 8 months. We concluded that our four focal hybrids PWK129S1, PWKB6, PWKAJ, and PWKDBA2 mice display three distinct trajectories of fertility: complete sterility (PWK129S1), complete fertility (pWkDBA2), and age-dependent fertility (PWKB6, PWKAJ). PWKWSB displays a fourth fertility trajectory that is likely influenced by its distinct *Prdm9^Dom3^* allele. This novel observation of age-dependent fertility reconciled our results with the literature and major breeding programs.

### Genomic patterns of subspecific origin implicate candidate HMS modifiers

The classical inbred strains in this study descend primarily from *M.m.domesiticus* ancestors, but have important contributions from *M.m.musculus*. The surprising fertility of PWKDBA2 hybrid males could be explained if the DBA2 genome had the greatest similarity to the PWK genome, compared to the genomes of 129S1, A/J, and B6. We compared each inbred strain to PWK and isolated areas of the genome where each inbred strain shared subspecific ancestry using publically available data [42]. We make no assumptions about whether the mode of action of these incompatibilities were due to underdominance at one locus, or by acting epistatically in conjunction with at least one additional locus. In addition, we also considered loci that shared ancestry regardless of subspecies identity, since PWK also has 5.72% *domesticus* ancestry. DBA2 did not share substantially more identity overall with PWK than the other three strains. DBA2 and PWK shared subspecific ancestry across 295.7 Mb of the genome (10.83%), similar to the shared ancestry between 129S1(10.30%), B6 (9.85%), and A/J (8.39%).

Nonetheless, we reasoned that the specific regions of the genome shared between PWK and specific classical strains but not others are good candidate locations for HMS modifier alleles. We focused on four specific contrasts based on the three patterns we observed in our experiments (**Figure 5, Table S2**). First, we searched for regions of the genome where only 129S1 differed in subspecific ancestry from PWK, reasoning that these regions may be enriched for incompatibility alleles unique to the 129S1 genetic background. We identified nine such regions (7.07 Mb) across six chromosomes. These regions contain 108 genes in total. Second, we searched for regions of the genome where A/J and B6 shared subspecific ancestry with each other but were different than DBA or 129S1. Regions where these strains also shared ancestry with PWK may harbor alleles that distinguish the age-dependent HMS in PWKAJ and PWKB6 males from the always-sterile PWK129S1 male. We found such seven regions (12.00 Mb) containing 146 genes. Regions in which A/J and B6 were alike but were different than the other three strains might harbor HMS alleles that explain the age-dependent effects. We found fourteen such regions (21.34 Mb) containing 111 genes. Finally, we searched for regions of the genome where only DBA2 shared subspecific ancestry with PWK. We reasoned that these regions might contain the critical modifier allele or alleles unique to DBA2 that rescue HMS in PWKDBA2 males. We discovered 44 such regions (87.65 Mb) across fifteen chromosomes. These loci contain a total of 834 genes. These candidate regions overlap several previously identified HMS QTL [21, 22, 36, 37] (black bars in Figure 5) and include genes that have been previously implicated in reproductive phenotypes.

**Figure 5:**
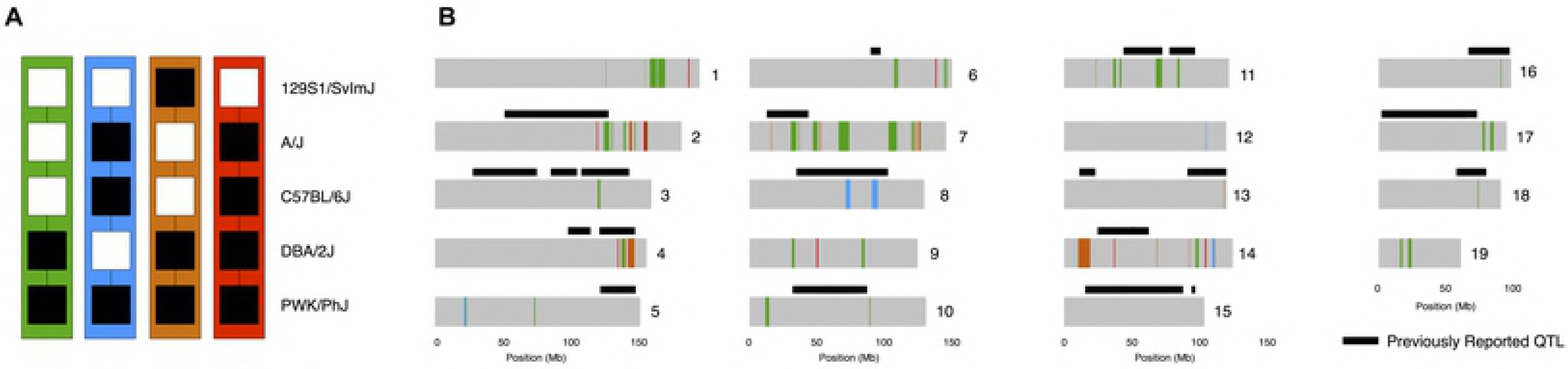
Subspecies haplotype sharing among inbred strains reveal candidate incompatibility loci. A) Contrasting patterns of subspecific ancestry among inbred strains that contribute to the observed phenotypes. Shared regions between PWK and DBA2 (green) may harbor alleles that rescue fertility in PWKDBA2 males. Regions shared between PWK, A/J, and B6 (blue) or regoins shared privately between B6 and A/J (orange) may associate with age-dependent HMS. Private incompatibilities between PWK and 129S1 (red) may underlie the complete sterility of PWK129S1 males. B) Chromosome plots (gray) map the haplotype sharing patterns from panel A. Black bars indicate the location of QTL identified in previous studies of HMS.

## Discussion

### Genetic architecture of HMS

Our results clearly show that HMS alleles segregate within the classical inbred strains, independent of the known major HMS loci. All the focal hybrids in this study share *Prdm9^Dom2/Msc^* genotype and carry the PWK Chr X. Chr Y is identical in the four classical inbred strain in our study. The three fertility trajectories we describe require at least two undiscovered autosomal factors. This supports a growing body of evidence that specific modifier alleles segregate in mice that rescue hybrids from male sterility. Several QTL have been associated previously with HMS phenotypes in both laboratory and wild-caught mice [35–38]. The ability to identify these modifiers has been hampered in most studies by segregating variation at the two major HMS loci. Our approach isolated the effects of modifiers that segregate within classical inbred strains, such that all of the phenotypic differences we see are due to the action of these modifiers. This advance is important because we have identified the specific strains that harbor modifier alleles. Furthermore, we have uncovered the importance of age to HMS and estimated the penetrance of HMS in PWKB6 hybrid males. With this prerequisite knowledge, our approach can be extended to QTL mapping to isolate and then identify specific modifier alleles.

The reciprocal hybrid males were all fertile, suggesting that the PWK Chr X was necessary for sterility in the focal hybrid males. However, the PWKWSB male sterility at 18 months shows that the age-dependent phenotypes are not dependent on the *Prdm9^Dom2^* allele, since WSB carries *Prdm9^Dom3^*. This supports previous findings that hybrids with *Prdm9^Dom3/Msc^* genotypes are categorically fertile but carry significantly reduced reproductive phenotypes in comparison to reciprocal hybrids [14, 36]. Furthermore, even the large-effect allele *Prdm9^Dom2^* is not sufficient for HMS, since the PWKDBA2 hybrid carries that allele and exhibits complete fertility.

The PWKDBA2 result allowed us to revaluate the hypothesis that mouse HMS is largely caused by diffuse genetic background differences throughout the genome that interact with *Prdm9*. If the DBA genome were more similar to PWK than the other classical inbred strains, which could explain the fertility of PWKDBA2 hybrids. However, our analysis showed that only a small fraction of the 129S1, A/J, B6, and DBA2 genomes showed *musculus* ancestry, and this fraction was roughly equal between the strains. We conclude that the dramatic phenotypic variation we see between these hybrids is most likely due to specific HMS modifier alleles segregating among those strains.

### Implications of age-dependent HMS

Steep allelic clines [11, 35] and reduced introgression on Chr X [10, 43] provide evidence that segregating HMS alleles are a partial reproductive barrier in the Central European mouse hybrid zone. If age-dependent HMS alleles segregate in the wild, they would allow certain hybrid males to escape infertility during a narrow window of life. Males exhibiting age-dependent HMS could hypothetically transmit HMS alleles to the next generation yet would have reduced relative fitness. Nonetheless, the window of fertility would attenuate the fitness cost of HMS and could promote the maintenance of the hybrid zone. Investigating this scenario will be difficult until we know the specific location of HMS modifiers.

Our results have immediate application to the increasingly popular Collaborative Cross and Diversity Outbred multi-parent mouse reference populations. The CC is a panel of recombinant inbred lines that are descended from eight inbred mouse strains including PWK, B6, A/J, and 129S1 [40, 41, 44] (DBA2 was not included), and DO is an outbred population descended from the same progenitors [45]. The CC breeding design necessarily depended on the focal hybrids derived from those strains. PWK129S1 hybrids were found to be sterile early in the breeding program [41] and did not contribute to CC lines. There was no specific observation of sterility in PWKAJ, PWKB6, or other hybrids offspring of PWK dams. However, most incipient CC lines (95%) stopped producing offspring during the inbreeding phase of the breeding program and were declared extinct. Nearly half (47%) of extinct lines contained sterile males [46], implicating HMS alleles segregating within the lines. Previously, we have concluded that epistasis involving 129S1 and PWK alleles were major drivers of male sterility and extinction. QTL mapping associated the PWK from a region on distal Chr X with male sterility [46]. However, each ancestral allele is only present in roughly an eighth of lines, and alleles from these two strains cannot explain the extinction rate entirely. HMS alleles harbored by B6 and A/J were segregating in the CC lines and could have had a substantial impact on extinction rate. Furthermore, the age-dependent sterility that we have described would have been difficult to detect, since fertility is incompletely penetrant and transient. According to the UNC Systems Genetics Core [47] that provides CC mice, fewer than 50% of males produce litters for nine different CC strains, suggesting that HMS modifier alleles were not eliminated during the breeding program and are relevant to current CC experiments as well as experimental crosses derived from CC strains.

### Mechanisms of age-dependent HMS

The molecular and cellular changes that confer fertility during such a narrow age range are unknown. Histological analysis revealed that meiosis was disrupted by 8 weeks of age and remained disrupted throughout life. Even transiently fertile PWKB6 males were distinctly different from reciprocal males in terms of their testis biology. On the other hand, the three focal hybrids had far more PMCs than the classic PWDB6 models of mouse HMS that have radically disorganized testes and few successful meioses [16, 18, 21]. These observations suggest that age-dependent HMS may be a threshold trait. If this hypothesis is true, PWKB6 and PWKAJ males at age 8 weeks had reproductive parameters just below the necessary threshold for fertility. Some factor that increased reproductive output around age 20 weeks was just enough to push some but not all of these males across the minimum required sperm count for fertility. 20-week sperm counts were elevated for all the focal and reciprocal hybrid males except for PWK129S1. This supports a threshold trait hypothesis, and suggests that slight increases in reproductive output in this age range are a normal part of male reproductive biology. The delayed onset of fertility could have been due to these normal changes on the background of abnormal meiosis. These critical differences might be driven by changes in testosterone (T) levels with age. At sexual maturity, testosterone initiates a cascade of events culminating in fertility including the induction of spermatogenesis [48]. However, seminal vesicle weights did not differ between sterile hybrids and their reciprocals (p = 0.155), and continued to increase with age (p ≤ 0.001); seminal vesicle weights are considered a suitable proxy for assessing serum T levels [49, 50]. We cannot explicitly rule out changes in serum or gonadal T over time as a driver of age-dependent HMS, but mice do not typically exhibit significant decreases in serum T in advanced age [51]. Nonetheless, future studies should measure hormone levels to test this hypothesis directly. Whatever biological factor drives the sperm increase at 20 weeks, PWK129S1 males uniquely show no response. Identifying the biology of this critical transition will point the way to the modifier allele harbored by the 129S1 mouse strain.

We expect both the delayed onset and premature cessation of sterility are linked to the depletion of PMCs observed at 8 weeks. However, it is unclear whether the proximal and genetic causes of the phenomena are the same. The reciprocal hybrid males showed no decline in reproductive parameters at 35 weeks. Furthermore, PWKWSB showed sterility at advanced age after a long period of fertility. Mitochondrial causes of age-related effects can be eliminated since all the focal hybrids inherited their mitochondria from a PWK dam, though mitochondrial-autosomal interactions cannot be ruled out.

An alternative mechanism for age-dependent decline is gene regulatory or epigenetic. *Prdm9* is a histone H3K4 methyltransferase and an essential epigenetic regulator of meiosis with the role of demarcating double-strand break sites where recombination will be initiated [28–31, 33]. This process is inherently sensitive to epigenetic changes. HMS modifier alleles could drive differential gene expression of target genes that directly interact with *Prdm9,* and that expression could be age dependent. A Prdm9-independent epigenetic mechanism is also possible. Faithful replication of epigenetic modifications is highly dependent on *trans-acting* genetic factors, in particular non-coding RNA (ncRNA, reviewed in [52]). Differentially expressed ncRNAs could alter the epigenomes of spermatogenic cells in a manner that renders them less competent to complete meiosis. Aberrant gene expression of Chr X has been repeatedly associated with HMS [24, 35, 37] and a compelling hypothesis is that HMS alleles impair both meiotic sex chromosome inactivation (MSCI) and postmeiotic sex chromatin repression (PSCR) [17, 25]. We saw a substantial number of seminiferous tubules with PMCs, even in sterile hybrid males. This suggests a majority of spermatogenic cells successfully underwent these processes. Detailed genomic studies across hybrids of different ages may identify key genes that mark the onset of sterility. Single-cell approaches may be especially appropriate given the divergent fates of individual spermatocytes.

In summary, we reported a wide range of male sterility in hybrid mice derived from the PWK strain. The differences between these hybrids were completely dependent on genetic background, which was invariant for genotypes previously associated with HMS at two major loci. We reported PWKDBA2 hybrid males as the only known mice to have these genotypes and yet completely escape HMS. Furthermore, we present the first known observation of HMS that onsets with age, and we characterized the unique fertility profiles associated with this age-dependent HMS. Taken together, these findings demonstrate both a novel phenotype and the classical inbred strains that harbor the relevant HMS modifier alleles. Identification of these modifiers will undoubtedly contribute to a growing body of work characterizing the genetic architecture of HMS in the mouse, and valuable insight into its mechanisms.

## Materials and Methods

### Mice

We generated F1 hybrid male mice by crossing PWK females to males of four inbred strains: 129S1, A/J, B6, and DBA2. Specific hybrids of this group will be collectively referred to as “focal hybrids” and individually referred to with the nomenclature “Dam Strain, Sire Strain” (i.e. PWK129S1 males are produced by crossing PWK females to 129S1 males). Focal hybrids (PWK129S1, PWKB6, PWKAJ, PWKDBA2) share *Prdm9^Dom2/Msc^* genotypes [15, 16, 18] and a are hemizygous for the PWK Chr X. We also bred the reciprocal hybrids (129S1PWK, AJPWK, B6PWK, and DBA2PWK) by crossing classical inbred strain females to PWK males. We also produced PWKWSB and WSBPWK hybrids, which carry *Prdm9^Dom3/Msc^*. All mice were fed soy-free Teklad mouse chow *ad-libitum*. All procedures involving animals were performed according to the Guide for the Care and Use of Laboratory Animals with approval by the Institutional Animal Care and Use Committee of North Carolina State University (NCSU) or the University of North Carolina at Chapel Hill (UNC-CH).

### Reproductive Phenotyping

Males were euthanized using carbon dioxide asphyxiation followed by cervical dislocation to confirm death. Weights for the carcass, testes, epididymides, and seminal vesicles were recorded. Sperm counts were collected from the right caudal epididymis after harvest. The whole epididymis was incubated in 500 μL of phosphate-buffered saline for at least 15 minutes at 37° C in an empty petri dish. Following incubation, the vas deferens and caput epididymis were removed and the cauda was snipped and incubated again for 15 minutes at 37° C. After the second incubation, sperm were extruded from the cauda using curved forceps. Once the sperm suspension was collected in a microcentrifuge tube, the petri dish was rinsed with additional PBS to collect remaining suspension, bringing the final suspension volume to 1 mL. Sperm was counted using a NucleoCounter SP-100 sperm cell counter (Chemometec). Left testes were fixed in Bouin’s solution overnight, and serially washed in 25%, 50%, and 70% EtOH. Testes were then embedded in paraffin wax, sectioned at 5 μm width, and stained according to a standard hematoxylin and eosin staining protocol. The number of seminiferous tubules containing post-meiotic cells was assessed in between 35 and 50 seminiferous tubules from each testis by counting the number of such tubules that meet this requirement in each image and averaging over the number of seminiferous tubules counted from each testis.

### Fertility Testing

We crossed 26 PWKB6, 6 PWKAJ, 3 PWKDBA2, and 6 B6PWK males to FVB females beginning between 5 and 8 weeks of age. Crosses were separated when FVB females were gravid or upon discovery of a litter, and continued only until 20 weeks of age when these mice were sacrificed for reproductive phenotyping. The age at which the male became fertile was calculated by subtracting 21 days from the litter’s birth date. Separately, we evaluated the fertility of 3 129S1PWK, 3 AJPWK, 2 B6PWK, 3 DBA2PWK, 3 WSBPWK, 3 PWK129S1, 5 PWKAJ, and 7 PWKDBA2, as controls. Lastly, we tested PWKB6 and PWKWSB males in a separate experiment conducted at UNC-CH. PWKB6 male mice (*n*=63) were continuously crossed to FVB females beginning at ages between 13-25 weeks. PWKWSB male mice (*n*=55) were continuously crossed to FVB females beginning at average age 21 weeks. Crosses were continually checked for litters until males ceased producing litters.

### Subspecific ancestry

Subspecific ancestry tracks for the 129S1, A/J, B6, DBA2, and PWK genomes were publically-available from the Mouse Phylogeny Viewer (http://msub.csbio.unc.edu/) [13, 42]. Each of the *domesticus* genomes was scanned for shared subspecific ancestry with PWK using the *Granges* package implemented through Bioconductor [53]. Shared genomic regions were then classified by whether were unique to an inbred strain (i.e. only 129S1 and PWK share common ancestry), or whether they shared subspecific ancestry with strains that, when crossed to PWK females, produce sterile hybrids at 8 weeks of age (129S1, A/J, and B6), fertile hybrids at 20 weeks of age (A/J, B6, DBA2), or hybrids that display age-dependent fertility with incomplete penetrance (A/J and B6). Genomic regions that shared PWK subspecific ancestry were then queried for known genes using the UCSC mouse genome table browser. We then queried these gene sets in the Mouse Genome Informatics database (Jackson Laboratory) for genes previously implicated in male reproductive system function.

### Statistical Analysis

We compared focal hybrids and their corresponding reciprocals using 2-way Analysis of Variance (ANOVA) and Tukey Honestly Significant Difference (HSD) test. We compared a specific focal hybrid and its reciprocal using Welch’s t-test, which accounts for unequal sample variances. We compared the four focal hybrids using 1-way ANOVA and HSD. We compared a specific focal hybrid and its reciprocal across ages using 2-way ANOVA and HSD. We compared a specific hybrid across ages using a 1-way ANOVA and HSD. We compared multiple focal hybrids across ages using 2-way ANOVA and HSD. We performed all statistical tests using R 3.3.1 software.

## Acknowledgements

We thank Aylor lab members Nicole Allard, Thomas Konneker, Connor McKenney, and Pei-Li Yao for helpful feedback and technical assistance, and the staff of the North Carolina State University Biological Resources Facility. We thank Jim Crowley for sharing his initial observation of premature sterility in PWKWSB hybrids. We thank Fernando Pardo-Manuel de Villena and Tim Bell at the University of North Carolina-Chapel Hill for supporting the breeding experiment of PWKB6 and PWKWSB hybrid males.

## Supporting Information Legends

**S1: Phenotype summary statistics.** Combined testes weight, sperm count, and percentage of seminiferous tubules containing postmeiotic cells (PMCs) across PWK-derived hybrids at each age point (Mean ± SE).

**S2: Genomic regions exhibiting specific patterns of subspecies haplotype sharing.** Chromosome, start sites, stop sites, and interval width expressed in megabases (Mb).

## References

1. Dobzhansky T. Genetics and the origin of species. New York: Columbia University Press; 1937.

2. Muller HJ. Isolating mechanisms, evolution, and temperature. Biol Symp. 1942;6:71–125.

3. Orr HA. The population genetics of speciation: The evolution of hybrid incompatibilities. Genetics. 1995;139:1805–13. doi: 10.1534/genetics.107.081810.

4. Boursot P, Auffray JC, Britton-Davidian J, Bonhomme F. The Evolution of House Mice. Annual Review of Ecology and Systematics. 1993;24(1993):119–52. doi: 10.1146/annurev.es.24.110193.001003.

5. Boursot P, Din W, Anand R, Darviche D, Dod B, VonDeimling F, et al. Origin and radiation of the house mouse: Mitochondrial DNA phylogeny. Journal of Evolutionary Biology. 1996;9(4):391–415. doi: 10.1046/j.1420-9101.1996.9040391.x.

6. Phifer-Rixey M, Nachman MW. Insights into mammalian biology from the wild house mouse Mus musculus. eLife. 2015;4:1–13. doi: 10.7554/eLife.05959.

7. Geraldes A, Basset P, Gibson B, Smith KL, Harr B, Yu HT, et al. Inferring the history of speciation in house mice from autosomal, X-linked, Y-linked and mitochondrial genes. Molecular Ecology. 2008;17(24):5349–63. doi: 10.1111/j.1365-294X.2008.04005.x.

8. Tucker PK, Sage RD, Warner J, Wilson aC, Eicher EM. Abrupt Cline for Sex Chromosomes in a Hybrid Zone between Two Species of Mice. Evolution. 1992;46(4):1146–63.

9. Teeter KC, Payseur BA, Harris LW, Bakewell MA, Thibodeau LM, O’Brien JE, et al. Genome-wide patterns of gene flow across a house mouse hybrid zone. Genome Research. 2008;18(1):67–76. doi: 10.1101/gr.6757907.

10. Janousek V, Wang L, Luzynski K, Dufková P, Vyskočilová MM, Nachman MW, et al. Genome-wide architecture of reproductive isolation in a naturally occurring hybrid zone between Mus musculus musculus and M. m. domesticus. Molecular Ecology. 2012;21:3032–47. doi: 10.1111/j.1365-294X.2012.05583.x.

11. Turner LM, Schwahn DJ, Harr B. Reduced male fertility is common but highly variable in form and severity in a natural house mouse hybrid zone. Evolution. 2012;66:443–58. doi: 10.1111/j.1558-5646.2011.01445.x.

12. Gregorová S, Forejt J. PWD/Ph and PWK/Ph inbred mouse strains of Mus m. musculus subspecies - A valuable resource of phenotypic variations and genomic polymorphisms. Folia Biologica. 2000;46(1):31–41.

13. Yang H, Wang JR, Didion JP, Buus RJ, Bell TA, Welsh CE, et al. Subspecific origin and haplotype diversity in the laboratory mouse. Nature Genetics. 2012;43(7):648–55. doi: 10.1038/ng.847.Subspecific.

14. Good JM, Handel MA, Nachman MW. Asymmetry and polymorphism of hybrid male sterility during the early stages of speciation in house mice. Evolution. 2008;62(1):50–65. doi: 10.1111/j.1558-5646.2007.00257.x.ASYMMETRY.

15. Mihola O, Trachtulec Z, Vlcek C, Schimenti JC, Forejt J. A mouse speciation gene encodes a meiotic histone H3 methyltransferase. Science (New York, NY). 2009;323(5912):373–5. doi: 10.1126/science.1163601.

16. Flachs P, Mihola O, Šimeček P, Gregorová S, Schimenti JC, Matsui Y, et al. Interallelic and Intergenic Incompatibilities of the Prdm9 (Hst1) Gene in Mouse Hybrid Sterility. PLoS Genetics. 2012;8(11). doi: 10.1371/journal.pgen.1003044.

17. Bhattacharyya T, Gregorova S, Mihola O, Anger M, Sebestova J, Denny P. Mechanistic basis of infertility of mouse intersubspecific hybrids. PNAS. 2013;110(6):E468–77. doi: https://doi.org/10.1073/pnas.1219126110.

18. Flachs P, Bhattacharyya T, Mihola O, Piálek J, Forejt J, Trachtulec Z. Prdm9 incompatibility controls oligospermia and delayed fertility but no selfish transmission in mouse intersubspecific hybrids. PLoS ONE. 2014;9(4). doi: 10.1371/journal.pone.0095806.

19. Davies AB, Hatton E, Altemose N, Hussin JG, Pratto F, Zhang G, et al. Re-engineering the zinc fingers of PRDM9 reverses hybrid sterility in mice. Nature. 2016;530(7589):171–6. doi: 10.1038/nature16931.

20. Hayashi K, Yoshida K, Matsui Y. A histone H3 methyltransferase controls epigenetic events required for meiotic prophase. Nature. 2005;438(November):374–8. doi: 10.1038/nature04112.

21. Bhattacharyya T, Reifova R, Gregorova S, Simecek P, Gergelits V, Mistrik M, et al. X Chromosome Control of Meiotic Chromosome Synapsis in Mouse Inter-Subspecific Hybrids. PLoS Genetics. 2014;10(2). doi: 10.1371/journal.pgen.1004088.

22. Dzur-Gejdosova M, Simecek P, Gregorova S, Bhattacharyya T, Forejt J. Dissecting the Genetic Architecture of F1 Hybrid Sterility in House Mice. Evolution. 2012;66:3321–35. doi: 10.5061/dryad.9cp1f.

23. Balcova M, Faltusova B, Gergelits V, Bhattacharyya T. Hybrid Sterility Locus on Chromosome X Controls Meiotic Recombination Rate in Mouse. PLoS Genetics. 2016;12(4). doi: doi:10.1371/journal.pgen.1005906.

24. Larson EL, Keeble S, Vanderpool D, Dean MD, Good JM. The Composite Regulatory Basis of the Large X-Effect in Mouse Speciation. Molecular Biology and Evolution. 2017;34(2):282–95. doi: 10.1093/molbev/msw243.

25. Campbell P, Good JM, Nachman MW. Meiotic sex chromosome inactivation is disrupted in sterile hybrid male house mice. Genetics. 2013;193(March):819–28. doi: 10.1534/genetics.112.148635.

26. Good JM, Giger T, Dean MD, Nachman MW. Widespread over-expression of the X chromosome in sterile F1 hybrid mice. PLoS genetics. 2010;6(9):e1001148–e. doi: 10.1371/journal.pgen.1001148.

27. Baudat F, Buard J, Grey C, Fledel-Alon A, Ober C, Przeworski M, et al. Prdm9 is a major determinant of meiotic recombination hotspots in humans and mice. Science. 2010;327(February):836–40. doi: 10.1044/2014.

28. Parvanov ED, Petkov PM, Paigen K. Prdm9 controls activation of mammalian recombination hotspots. Science. 2010;327(February):835-.

29. Baker CL, Kajita S, Walker M, Saxl RL, Raghupathy N, Choi K, et al. PRDM9 Drives Evolutionary Erosion of Hotspots in Mus musculus through Haplotype-Specific Initiation of Meiotic Recombination. PLoS genetics. 2015;11(1):e1004916–e. doi: 10.1371/journal.pgen.1004916.

30. Baker CL, Walker M, Kajita S, Petkov PM, Paigen K. PRDM9 binding organizes hotspot nucleosomes and limits Holliday junction migration. Genome research. 2014;24(5):724–32. doi: 10.1101/gr.170167.113.

31. Brick K, Smagulova F, Khil P, Camerini-Otero RD, Petukhova GV. Genetic recombination is directed away from functional genomic elements in mice. Nature. 2012;485(7400):642–5. doi: 10.1038/nature11089.

32. Walker M, Billings T, Baker CL, Powers N, Tian H, Saxl RL, et al. Affinity-seq detects genome-wide PRDM9 binding sites and reveals the impact of prior chromatin modifications on mammalian recombination hotspot usage. Epigenetics & Chromatin. 2015;8:13. doi: 10.1186/s13072-015-0024-6.

33. Grey C, Barthès P, Friec G, Langa F, Baudat F, de Massy B. Mouse Prdm9 DNA-binding specificity determines sites of histone H3 lysine 4 trimethylation for initiation of meiotic recombination. PLoS Biology. 2011;9(10):1–9. doi: 10.1371/journal.pbio.1001176.

34. Gregorova S, Gergelits V, Chvatalova I, Bhattacharyya T, Valiskova B, Fotopulosova V, et al. Modulation of Prdm9-controlled meiotic chromosome asynapsis overrides hybrid sterility in mice. Elife. 2018;7. doi: 10.7554/eLife.34282. PubMed PMID: 29537370.

35. Turner LM, Harr B. Genome-wide mapping in a house mouse hybrid zone reveals hybrid sterility loci and Dobzhansky-Muller interactions. eLife. 2014;3(2002):1–25. doi: 10.7554/eLife.02504.

36. White MA, Steffy B, Wiltshire T, Payseur BA. Genetic dissection of a key reproductive barrier between nascent species of house mice. Genetics. 2011;189(1):289–304. doi: 10.1534/genetics.111.129171.

37. Turner LM, White MA, Tautz D, Payseur BA. Genomic networks of hybrid sterility. PLoS genetics. 2014;10(2):e1004162–e. doi: 10.1371/journal.pgen.1004162.

38. Larson EL, Vanderpool D, Sarver BAJ, Callahan C, Keeble S, Provencio LP, et al. The Evolution of Polymorphic Hybrid Incompatibilities in House Mice. Genetics. 2018. doi: 10.1534/genetics.118.300840.

39. Consortium CC. The Genome Architecture of the Collaborative Cross Mouse Genetic Reference Population. Genetics. 2012;190(2):389–401. doi: 10.1534/genetics.111.132639.

40. Churchill GA, Airey DC, Allayee H, Angel JM, Attie AD, Beatty J, et al. The Collaborative Cross, a community resource for the genetic analysis of complex traits. Nature genetics. 2004;36(11):1133–7. doi: 10.1038/ng1104-1133.

41. Chesler EJ, Miller DR, Branstetter LR, Galloway LD, Jackson BL, Philip VM, et al. The Collaborative Cross at Oak Ridge National Laboratory: developing a powerful resource for systems genetics. Mammalian Genome. 2008;19(6):382–9. doi: 10.1007/s00335-008-9135-8.

42. Wang JR, de Villena FP, McMillan L. Comparative analysis and visualization of multiple collinear genomes. BMC Bioinformatics. 2012;13 Suppl 3:S13. doi: 10.1186/1471-2105-13-S3-S13. PubMed PMID: 22536897; PubMed Central PMCID: PMC3311102.

43. Payseur BA, Krenz JG, Nachman MW. Differential patterns of introgression across the X chromosome in a hybrid zone between two species of house mice. Evolution; international journal of organic evolution. 2004;58(9):2064–78. doi: 10.1554/03-738.

44. Aylor DL, Valdar W, Foulds-mathes W, Buus RJ, Verdugo RA, Baric RS, et al. Genetic analysis of complex traits in the emerging Collaborative Cross. 2011;(Churchill 2007):1213–22. doi: 10.1101/gr.111310.110.mentally.

45. Churchill GA, Gatti DM, Munger SC, Svenson KL. The Diversity Outbred mouse population. Mamm Genome. 2012;23(9–10):713–8. doi: 10.1007/s00335-012-9414-2. PubMed PMID: 22892839; PubMed Central PMCID: PMC3524832.

46. Shorter JR, Odet F, Aylor DL, Pan W, Kao CY, Fu CP, et al. Male Infertility Is Responsible for Nearly Half of the Extinction Observed in the Mouse Collaborative Cross. Genetics. 2017;206(2):557–72. doi: 10.1534/genetics.116.199596. PubMed PMID: 28592496.

47. Welsh CE, McMillan L. Accelerating the inbreeding of multi-parental recombinant inbred lines generated by sibling matings. G3 (Bethesda). 2012;2(2):191–8. doi: 10.1534/g3.111.001784. PubMed PMID: 22384397; PubMed Central PMCID: PMC3284326.

48. Singh J, Oneill C, Handelsman DJ. Induction of spermatogenesis by androgens in gonadotropin-deficient (hpg) mice. Endocrinology. 1995;136(12):5311–21. doi: 10.1210/en.136.12.5311.

49. Bartke A, Shire JGM. Differences between mouse strains in testicular cholesterol levels and androgen target organs. Journal of Endocrinology. 1972;55(1):173–84. doi: 10.1677/joe.0.0550173.

50. Bartke A. Increased sensitivity of seminal vesicles to testosterone in a mouse strain with low plasma testosterone levels. Journal of Endocrinology. 1974;60(1):145–8. doi: 10.1677/joe.0.0600145.

51. Eleftheriou BE, Lucas LA. Age-related changes in testes, seminal vesicles and plasma testosterone levels in male mice. Gerontologia. 1974;20(4):231–8.

52. Margueron R, Reinberg D. Chromatin structure and the inheritance of epigenetic information. Nature Reviews Genetics. 2010;11(4):285–96. doi: 10.1038/nrg2752.

53. Lawrence M, Huber W, Pages H, Aboyoun P, Carlson M, Gentleman R, et al. Software for computing and annotating genomic ranges. PLoS Comput Biol. 2013;9(8):e1003118. doi: 10.1371/journal.pcbi.1003118. PubMed PMID: 23950696; PubMed Central PMCID: PMC3738458.

